# Rapid updating of ecological information in wild Guinea baboons

**DOI:** 10.1101/2025.09.17.676797

**Authors:** Lisa Ohrndorf, Roger Mundry, Dietmar Zinner, Julia Fischer

## Abstract

Knowing where to find what kind of food when is essential for survival and reproduction. Recent links between feeding ecology and brain size have revived interest in foraging cognition as a core component in cognitive evolution. We investigated whether free-ranging Guinea baboons (*Papio papio*) keep track of the spatio-temporal availability of a preferred food resource, the Natal orange (*Strychnos spinosa*). A natural experiment, a massive wildfire that destroyed most oranges in 2019, allowed us to disentangle habitual routines from goal-directed behaviour in the baboons. The animals’ space-use patterns shifted significantly following the wildfire. After one single visit, they stopped going to the area during the period when Natal oranges would normally be available. In the following year, they resumed regular space-use patterns. We also assessed travel linearity, speed, and departure time from sleeping sites towards the target area with and without ripe fruit. Guinea baboons travelled more linearly, 14% faster, and ~40 minutes earlier towards the target area when *S. spinosa* was in season compared to when it was not. In conclusion, Guinea baboons rapidly updated their knowledge of high-value food availability and adjusted their space-use accordingly. These results underscore their capacity for flexible decision-making and goal-directed foraging.

## Introduction

Knowing ‘where’ to find ‘what’, ‘when’ it is available, and how to move between specific locations is essential to maximise foraging outcomes while simultaneously minimising associated costs. Navigational capacities can thus have important implications for the survival and reproduction of animals. Recently established links between feeding ecology and brain size have revived interest in foraging cognition as a central driver of cognitive evolution [1–4]. One way in which animals may achieve efficient foraging is to memorise the locations of valuable resources and travel directly towards these out-of-sight goals [5,6]. Such ‘goal-directedness’ is often inferred from observed movement patterns such as increased travel linearity, faster approaches, or earlier departures towards high-value feeding sites [6–8].

Several studies across taxa have highlighted animals’ abilities to optimise travel paths towards high-value resources, frequently demonstrating spatial memory. Insects such as fruit flies (*Drosophila melanogaster*), honeybees (*Apis mellifera*), and bumblebees (*Bombus* spp.) demonstrated robust spatial learning abilities and efficient trap-lining between food resources [9–12]. Birds have long served as key models for studying the navigational capacity of animals [13], from homing pigeons (*Columba livia*) to a wide range of other species such as honey buzzards (*Pernis ptilorhynchus)* [14], cuckoos (*Cuculus canoris*) [15] and rufous hummingbirds (*Rufus selasphorus*) [16], with research highlighting the complementary use of cue-based and memory-based navigation [17,18]. Among primates, Western gorillas (*Gorilla gorilla*), chacma baboons (*Papio ursinus*), and Western chimpanzees (*Pan troglodytes verus*) show space-use patterns consistent with goal-directed foraging, including linear travel, selective bypassing of low-quality food resources, and earlier departure towards valuable feeding trees [6,8,19].

Yet, observed space-use patterns alone are often insufficient to differentiate between habitual and goal-directed behaviour. Linearity, for example, may arise not only from deliberate targeting of a known resource but also from landscape constraints or simple rules such as avoiding backtracking [20]. Similarly, faster travel or earlier departures could reflect fixed routines rather than flexible, goal-directed decision-making [20]. This ambiguity is further compounded by differences in how ‘goal-directedness’ is defined across research domains. In the study of spatial cognition, the term ‘goal-directedness’ typically refers to efficient travel towards out-of-sight resources, whereas in the domain of associative learning it denotes actions that are flexibly updated when outcomes change, in contrast to habitual routines that persist irrespective of current payoffs [21,22]. In a foraging context, both approaches likely capture complementary aspects of the same functional principle: animals adjust their space-use to maximise rewards and minimise costs. Goal-directedness may thus be expressed both as efficient, directed travel towards a localised food resource, and as outcome-sensitive adjustments of space-use when payoffs shift. Here, we use this overlap to interpret observed movement patterns as indicators of flexible, reward-sensitive decision-making and goal-directed foraging.

In the present study, we analysed Guinea baboon (*Papio papio*) movement patterns towards a valuable, localised, and temporally available food resource, the Natal orange (*Strychnos spinosa*). These orange-sized fruits are enclosed in a firm shell and contain tightly packed seeds surrounded by a layer of fleshy, edible pulp. Of particular interest was how the baboons’ space-use patterns changed when this food resource was unexpectedly not available during the typical season. At our long-term field site, Centre de Recherche du Primatologie Simenti in the Niokolo Koba National Park in south-east Senegal [23] (Figure 1), the Natal orange is one of the few feeding tree species that provide fleshy fruit outside the rainy season [24]. The trees typically bear ripe fruit in the dry season from January to March.

**Figure 1.**
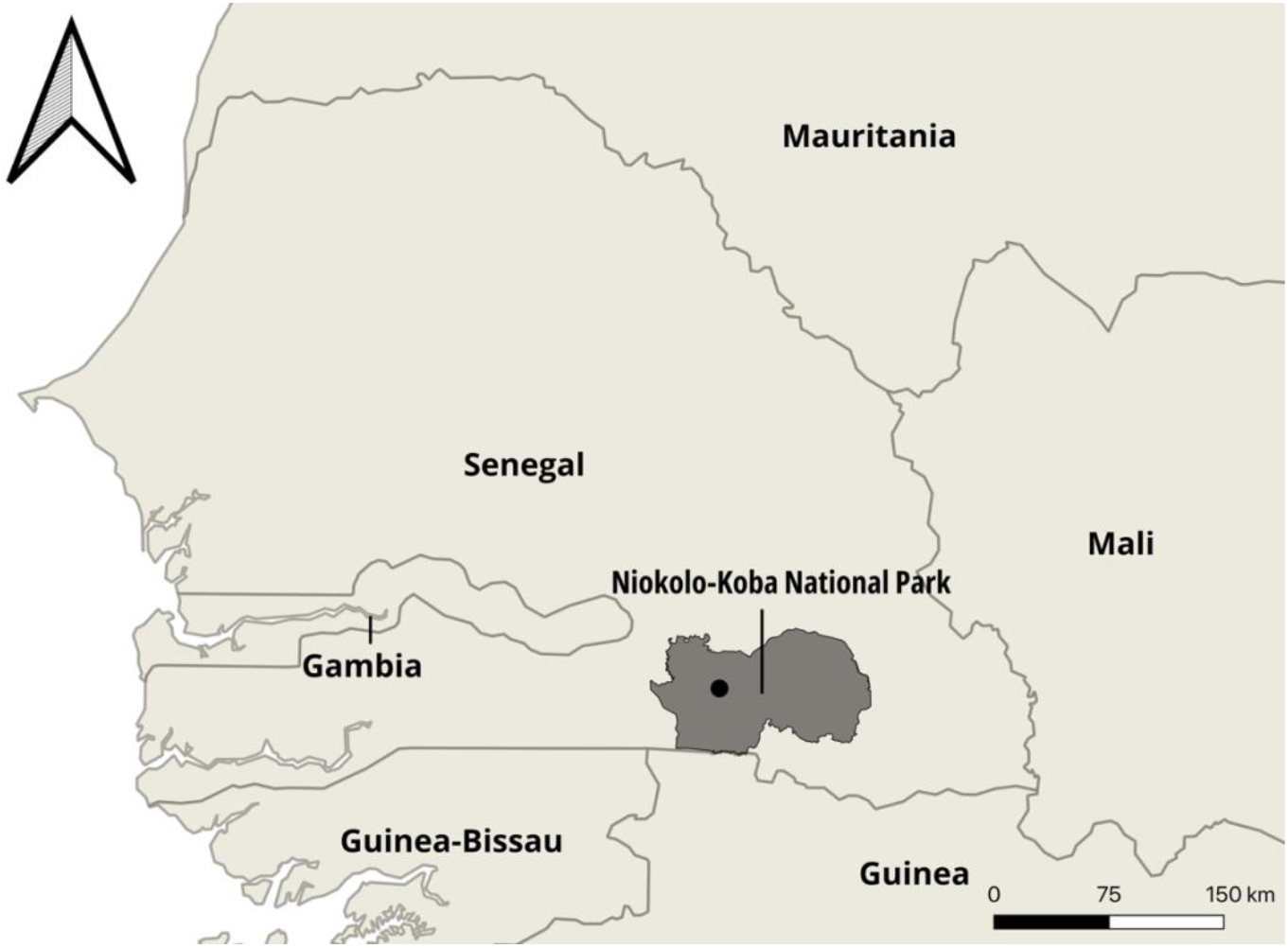
Location of the field site of the CRP Simenti (black dot) within the Niokolo-Koba National Park (PNNK).

To shed light on the ‘cognitive processing architecture’ [25] underlying the animals’ space-use, that is, the distinction between habitual and goal-directed decision-making, we made use of a ‘natural experiment’ that occurred in the dry season of 2019. In late January 2019, the National Park experienced extensive bushfires, during which most of the fruits of *S. spinosa* burned. We assessed whether the baboons showed a shift in their space-use patterns in 2019 compared to two years prior and two years after the destruction of fruits by bushfires. If the baboons’ movement patterns are predominantly habitual, we would expect them to follow their usual routines also in 2019. However, if they learnt from inspecting the area that the fruit was not available (and would also not become available later), they should stop visiting the area after a few visits.

We further assessed the linearity and speed of travel, as well as the time of departure from sleeping sites towards the target area when ripe fruits were available, compared to when they were not. If the baboons mentally represent the availability of the food resource, they should depart from the sleeping trees earlier and travel more linearly and faster when Natal oranges are available.

## Material and Methods

### Study site

Data collection took place at the Centre de Recherche de Primatologie (CRP), Simenti (13°01’34’’ N, 13°17’41’’ W) in the Niokolo Koba National Park (PNNK) in South-East Senegal (Figure 1). The study site is part of the Sudanian and Sahelo-Sudanian climatic zone with pronounced seasonality and high variability of rainfall among seasons [26,27]. The rainy season typically lasts from June to October, with May and mid-October constituting transitional periods with little and very variable rainfall [26]. From 2017 to 2022, the average annual precipitation was 950 mm. The vegetation represents a mosaic of grasslands, wooded savannahs, and gallery forests along streams or larger water bodies [26,28]. Several habituated parties of Guinea baboons live at the study site, which have been followed extensively by researchers since 2007 [23].

### Study species

Guinea baboons occupy a variety of habitats, from moist forests and mangroves in Guinea to semi-deserts in Mauritania, and are likely exposed to considerable seasonal variation in precipitation across their range [24,29,30]. They are predominantly terrestrial and live in large groups comprising 20 to more than 300 individuals with a nested multi-level social organisation and female-biased dispersal [23, 31,32]. The first level of social organisation is the one-male unit (OMU) consisting of one adult male, one to several females, their dependent offspring, and occasionally secondary males [33]. Several of these OMUs form parties which then, together with other parties, form gangs [23]. The home ranges of parties in the Niokolo-Koba National Park span 20 to 50 km^2^ on average [24,34] with considerable overlap between them [35]. Like other baboon species, Guinea baboons are considered eclectic omnivores with a high proportion of fruit in their diet [24,34]. At the study site, the Natal orange (*S. spinosa*) is one of the few feeding tree species that provide fleshy fruit outside of the rainy season [24]. Trees of this species predominantly occur in the northern part of the home range of our study parties. It typically bears ripe fruit from January to March, which the baboons feed on preferentially during this time (personal observation).

### Data collection

#### Target area

To delineate the area in which *S. spinosa* occurs at a high density, we established 10 parallel transects in a North-South direction of roughly 4 km length with 500 m between transects. We noted the number and locations of all *S. spinosa* trees within 5 m of the transect line. We ran a kernel density estimation in QGIS 3.30 using the heatmap function with a quartic kernel shape and a radius/bandwidth of 500 m. We used the 95% contour level to delineate the “target area” (Figure 2a).

**Figure 2.**
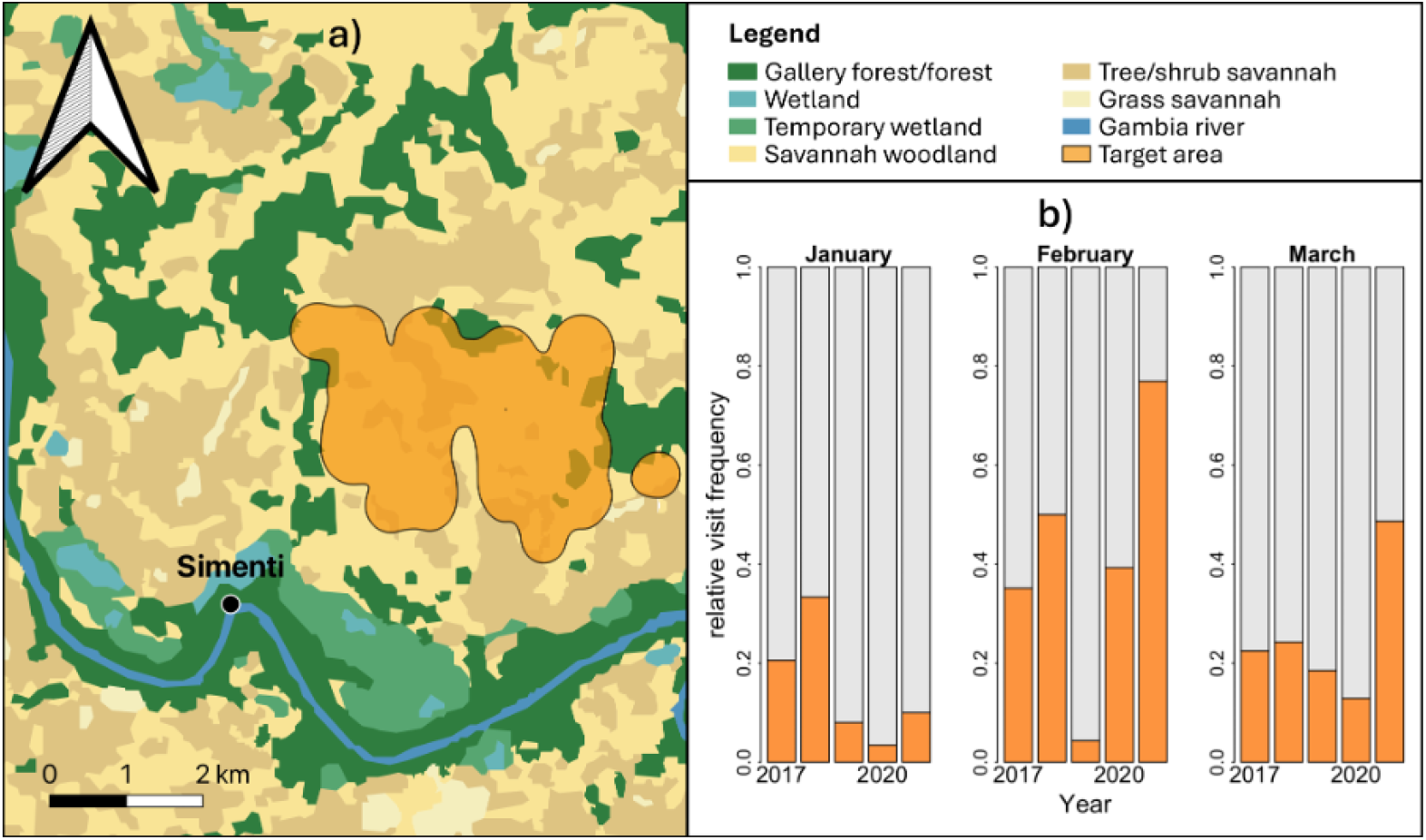
Guinea baboons stop visiting the target area after the destruction of ripe fruit by bushfires in 2019. a) shows the location of the “Target area” representing the occurrence of *S. spinosa* trees at high densities (277 trees per hectare, delineated using kernel density estimation (KDE) at the 95% contour level) and b) depicts the relative frequency of visits to the target area in January, February, and March from 2017 to 2021. Orange sections of the bars indicate the proportion of days on which at least one waypoint from the study party fell within the target area, relative to the total number of observation days.

#### Season of S. spinosa

We established the time at which ripe fruits of *S. spinosa* were most abundant using phenological data from 2013 to 2014 and from 2022 to 2023. Although fruits are already available in December, they only start to ripen in January, and ripe fruits are most abundant in February, with some lasting until March. We therefore established the “season” to last from January to March every year. We classified all remaining months as “non-season”. We also classified February and March of 2019 as a non-season, because the majority of ripe fruits had burned.

#### GPS data

Locational data of Guinea baboons were collected through direct observation during half-day follows (typically from 07:00 to 13:00h) of the baboon parties using Garmin handheld GPS devices (GARMIN GPSmap 62-66s) from April 2014 to March 2023. During the follows, waypoints were taken at 30-minute intervals. The initial waypoint for a given day was taken upon first contact with a specific party, and, accordingly, the last waypoint was taken upon the end of baboon contact.

#### Linearity and speed of travel

To assess the parameters of space-use, we used a subset of the overall GPS data set. We only considered days on which the baboons moved towards the target area. We omitted tracks that consisted of two waypoints or fewer before the baboons entered the target area. We then calculated a simple linearity index by dividing the Euclidean distance from the first point of a given day to the first point of that day that was within the target area by the sum of the distances between successive waypoints leading up to the target area (Figure 3). We calculated the mean travel speed by dividing the sum of the distances between successive waypoints leading up to the target area by the sum of the time lags between the respective waypoints.

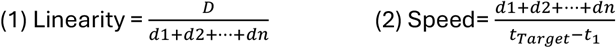

**Figure 3.**
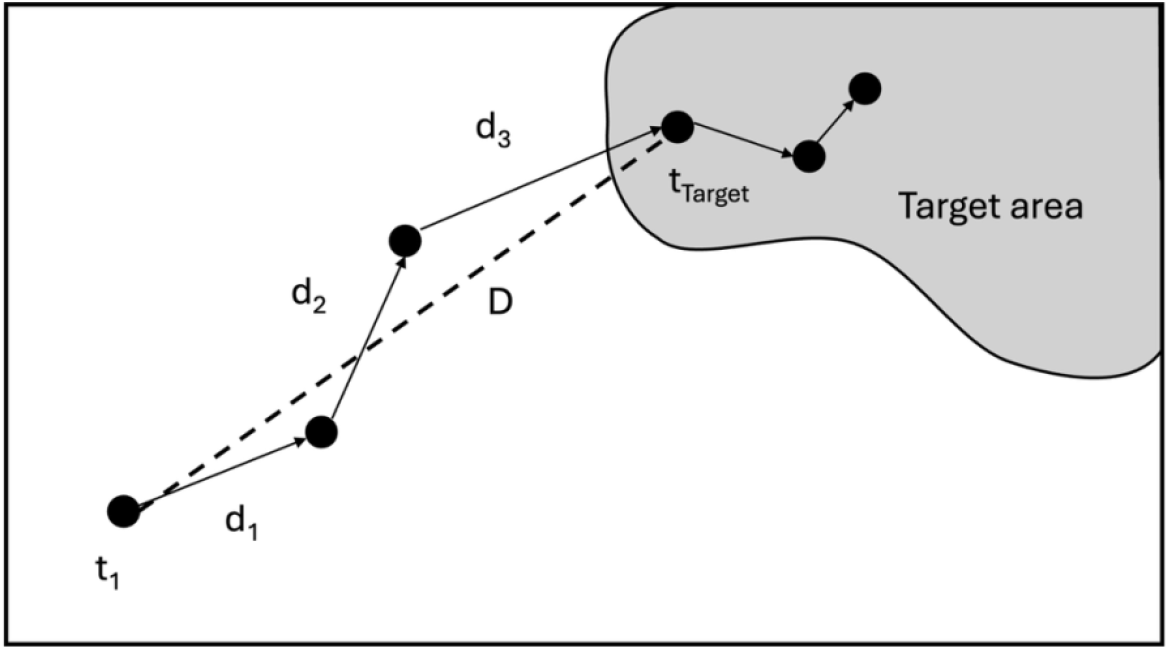
Schematic depiction of calculations for linearity and speed of travel. D = Euclidean distance travelled, d_n_ = distance between successive waypoints, t_1_ = time of the day’s first waypoint (at first contact), t_n_= time of waypoint n, t_Target_= time of first waypoint in target area.

We took waypoints only every 30 minutes following our general routine, whereas most other comparable studies chose shorter sampling intervals of 1 min [36], 2 min [37], or 5 min [6,19,38]. However, 10-minute [39–41] or 15-minute intervals [42,43] have also been used. Because of the coarse sampling, we likely underestimated the actual distance walked. Sennhenn-Reulen et al. [44] found an underestimation of ca. 25% of the actual travel distance at a 30-min sampling interval in the same study population. As a result, our linearity values are likely overestimated relative to the true linearity of baboon travel, and travel speed is likely underestimated. Nonetheless, as our analyses are based on internal comparisons and modelling within this dataset, the observed patterns and relative effects are robust to the sampling interval.

#### Time of departure from sleeping sites

We defined the time of departure from the sleeping sites as the moment when the baboons left a radius of 300 m around the sleeping site. In our analysis, we only included days on which we found the baboons at their sleeping sites. We then calculated the relative time of departure from the sleeping site as the time difference between sunrise and the time the baboons left the sleeping site. We derived times of sunrise for the study area using the package *suncalc* in R.

#### Temperature

Since the departure of the baboons from the sleeping site and their travelling may be affected by temperature (e.g., later departure due to sunbathing after colder nights), we controlled for the effects of temperature on the time of departure from the sleeping sites. We noted the daily minimum and maximum temperature using a digital thermometer (Eurochrom ETH 5500 Thermometer/Hygrometer) positioned outdoors at the campsite but protected from direct sunlight and precipitation.

### Statistical analyses

#### Natural experiment

We assessed changes in space-use patterns in response to pronounced bushfires in the study area in 2019. We calculated the number of days between January and March from 2017 to 2021 during which at least one waypoint for a study party fell into the target area. We then compared the relative frequency of visits to the target area across these years.

#### Linearity of travel (Model 1)

To estimate how linearity differed between season and non-season, we fitted a generalized linear mixed model (GLMM) [45] with a beta error structure and logit link function [46] using the ‘glmmTMB’ package version 1.1.7 [47] in R version 4.3.1 [48]. We used the linearity index as a response and season (yes/no) as a fixed effect. As we had repeated observations of several different study parties, we included party ID as a random intercept effect. We also included the random slope of season (yes/no) in the model to keep type I error rates at a nominal level of 0.05 [49,50]. Initially, we also included estimates of the correlations between random intercept and slope in the model. However, model fit did not decrease when removing the correlation from the model (log-likelihoods: full model including correlation: 644.33, df=6; full model without correlation: 643.37, df=5). We therefore continued the analysis with the model lacking the correlation parameter (Table 1).

**Table 1:**
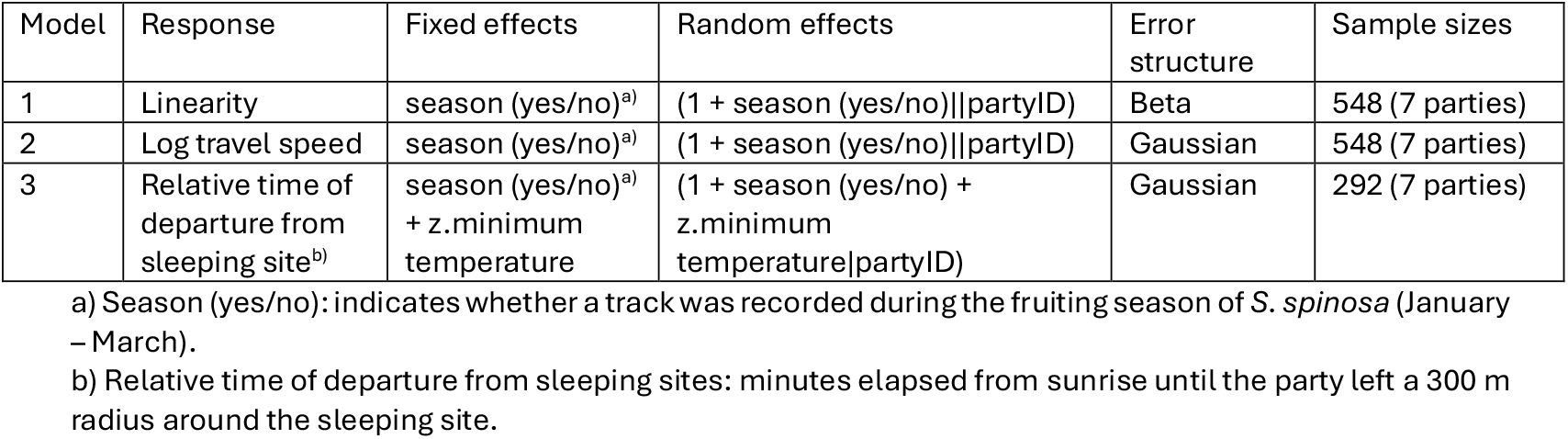
Overview of the fitted models.

#### Travel speed and relative time of departure from sleeping sites (Models 2 and 3)

For travel speed and the relative time of departure from the sleeping site, we fitted two linear mixed models (LMM) [45] with a Gaussian error structure and identity link function [46] using the function ‘lmer’ of the ‘lme4’ package version 1.1–33.1 [51]. We log-transformed travel speed to increase the probability of model convergence. We included log speed as the response variable and season (yes/no) as a fixed effect in model 2. For model 3, we included relative time of departure from sleeping sites as the response variable and season (yes/no) as a fixed effect. To control for the effects of temperature on time of departure from sleeping sites, we z-transformed the minimum temperature recorded on a given day to a mean of 0 and a standard deviation of 1 and added it as a fixed effect in model 3. As for model 1, we included party ID as a random intercept effect in both models. We further included the random slope of season (yes/no) in the models to keep type I error rates at a nominal level of 0.05 [49, 50]. Model 3 additionally included the random slope of minimum temperature. As for model 1, we initially included estimates of the correlation between random intercepts and slopes in model 2 and model 3. For model 2, model fit did not decrease when removing the correlation (log-likelihoods: full model including correlation: −306.40, df=6; full model without correlation: −306.67, df=5), so we proceeded with the analysis with the model lacking correlations. Model 3 included correlations between random intercept and slopes (Table 1). We used the function ‘lmer’ from the package ‘lmertest’ (version 3.1-3) [52] to test the effect of our fixed effects employing the Satterthwaite approximation [53].

#### General considerations

For model 1, we checked whether the assumption of the absence of overdispersion was met. With a dispersion parameter of 1.18, the response was not overdispersed given the model. After fitting models 2 and 3, we checked whether the assumption of homogeneity and normal distribution of residuals was fulfilled by visually inspecting a QQ-plot [54] of residuals and residuals plotted against fitted values [55]. These showed that assumptions were met. For all models, we assessed model stability by dropping each party from the data one at a time and comparing the estimates from models fitted to these subsets of data to those derived from the respective model for the full dataset. All models were of good stability (see results). We obtained confidence intervals of model estimates and fitted values using parametric bootstraps (N=1000; functions ‘simulate’ and ‘bootMer’ of the packages ‘glmmTMB’ and ‘lme4’, respectively).

## Results

### Natural experiment

For the period between January and March from 2017 to 2021, we collected a total of 471 tracks of seven study parties (N tracks = 111 in 2017, 91 in 2018, 75 in 2019, 88 in 2020, and 106 in 2021). During this time, the Guinea baboon parties visited the target area on 137 days (29 in 2017, 33 in 2018, 8 in 2019, 16 in 2020, 51 in 2021). Visit frequencies in January ranged from 1 to 8 across all years. In February, when ripe fruits of *S. spinosa* were typically most abundant, baboons visited the area once in 2019 compared to 11-30 times in the other study years. In March of 2019, the baboons visited the area five times, with visit frequencies ranging from 4 to 18 in the other study years (Figure 2b).

### Linearity and speed of travel (Models 1 and 2)

From April 2014 to March 2023, we collected 2938 tracks, comprising 31,028 location points, for seven study parties. For 548 tracks, at least one waypoint fell into the target area. 244 (44.5%) of these tracks were collected during the Natal oranges’ season, whereas 304 (55.5%) were collected outside the season. Tracks consisted of more than three waypoints, with an average of six waypoints (median, range 3-14) per track. The average linearity of travel paths was 0.89 ± 0.13 (mean, SD, range 0.29-1.00), where values of 1 signify completely linear travel trajectories and values of 0 indicate “meandering” travel.

Linearity indices were above 0.8 for 84%, and above 0.9 for 61% of all considered travel paths towards the target area. The linearity of travel towards the target area was higher during the Natal oranges’ season compared to the non-season (Table 2; Figure 4a). The Guinea baboon parties travelled at an average speed of 1.12 km/h ± 0.48 (mean, SD), ranging from 0.16 to 3.65 km/h. The baboons travelled faster towards the target area when Natal oranges were in season vs. when they were not. More specifically, they travelled 14% faster towards the target area when Natal oranges were in season compared to when they were not (Table 2, Figure 4b).

**Table 2:**
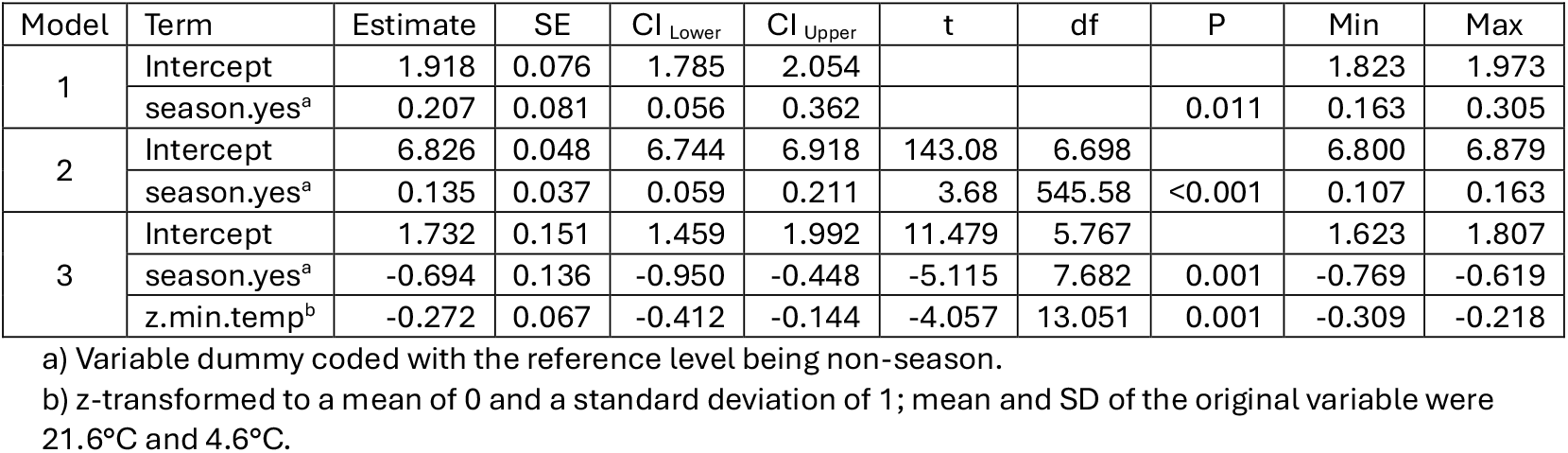
Model results on linearity (Model 1), speed (Model 2) and time of departure (Model 3); estimates, standard errors, confidence intervals, significance test, and range of estimates derived from dropping each party one at a time.

| Model | Term | Estimate | SE | CI<sub>Lower</sub> | CI<sub>Upper</sub> | t | df | P | Min | Max |
| --- | --- | --- | --- | --- | --- | --- | --- | --- | --- | --- |
| 1 | Intercept | 1.918 | 0.076 | 1.785 | 2.054 |  |  |  | 1.823 | 1.973 |
|  | season.yes<sup>a</sup> | 0.207 | 0.081 | 0.056 | 0.362 |  |  | 0.011 | 0.163 | 0.305 |
| 2 | Intercept | 6.826 | 0.048 | 6.744 | 6.918 | 143.08 | 6.698 |  | 6.800 | 6.879 |
|  | season.yes<sup>a</sup> | 0.135 | 0.037 | 0.059 | 0.211 | 3.68 | 545.58 | <0.001 | 0.107 | 0.163 |
| 3 | Intercept | 1.732 | 0.151 | 1.459 | 1.992 | 11.479 | 5.767 |  | 1.623 | 1.807 |
|  | season.yes<sup>a</sup> | -0.694 | 0.136 | -0.950 | -0.448 | -5.115 | 7.682 | 0.001 | -0.769 | -0.619 |
|  | z.min.temp<sup>b</sup> | -0.272 | 0.067 | -0.412 | -0.144 | -4.057 | 13.051 | 0.001 | -0.309 | -0.218 |

a) Variable dummy coded with the reference level being non-season.
b) z-transformed to a mean of 0 and a standard deviation of 1; mean and SD of the original variable were 21.6°C and 4.6°C.

**Figure 4.**
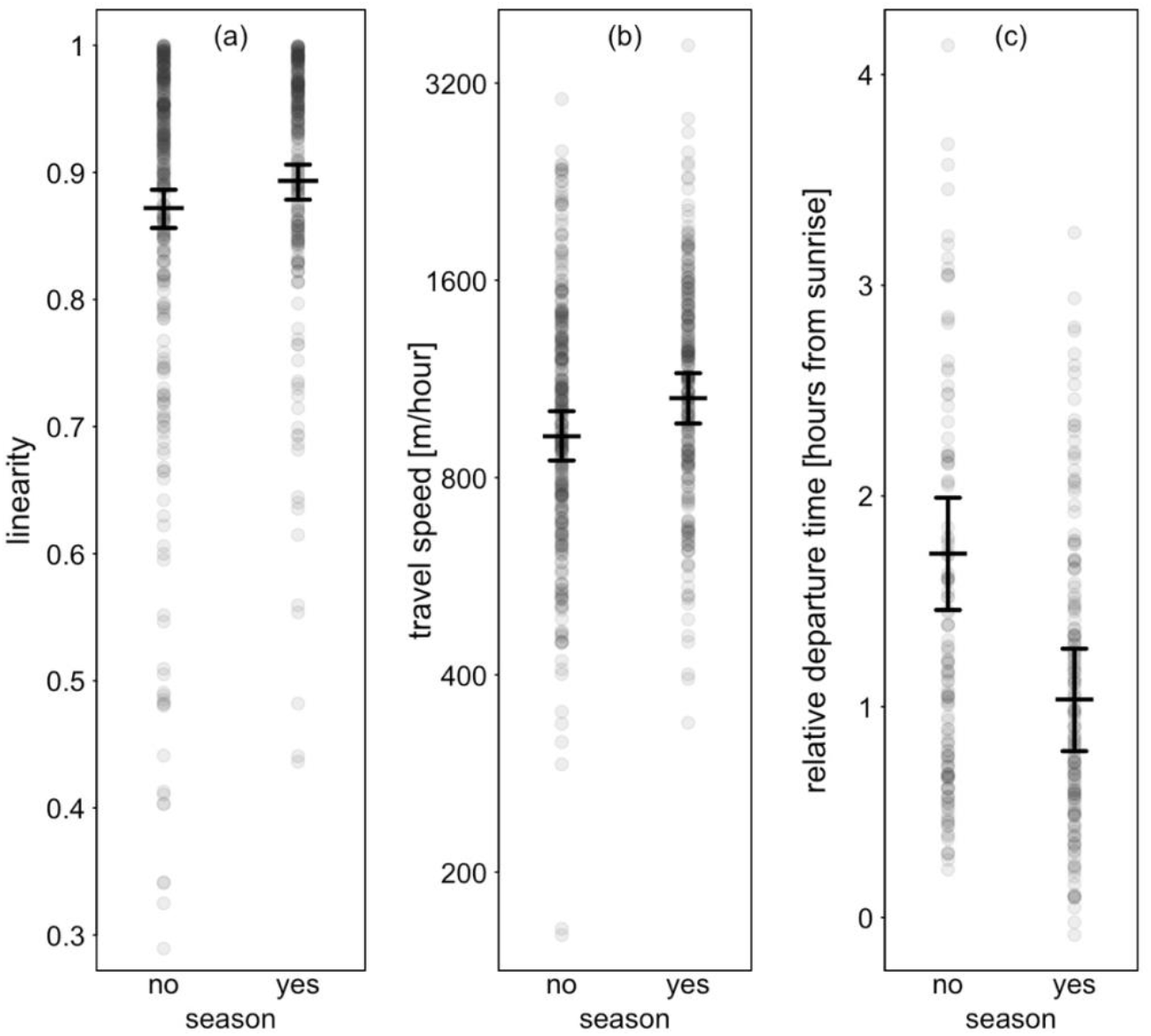
Guinea baboons travel more linearly, faster and leave earlier towards the target area when ripe fruits are available. Panels show (a) linearity, (b) travel speed, and (c) relative time of departure from sleeping sites towards the target area in seasons vs. non-seasons. Black lines depict fitted means and 95% confidence intervals.

Data on both the time of departure from sleeping sites and the minimum temperature were available for 292 tracks. Of these, 162 (55.5%) were collected during the season and 130 (44.5%) outside of the season. In the mornings, the baboons left their sleeping sites at 08:10h ± 50 min on average (mean, SD). On average, parties left their sleeping sites 1h 14 min ± 48 min (mean, SD) after sunrise. The minimum temperature for the study days ranged from 11°C to 31.8°C with an average of 21.6°C ± 4.6 (mean, SD). The Guinea baboons left their sleeping sites ca. 41 min earlier when travelling towards the target area when fruits of *S. spinosa* were in season vs. when they were not (Table 2; Figure 4c). The Guinea baboons left their sleeping sites significantly later when minimum temperatures were lower (Table 2).

## Discussion

Our findings revealed a substantial shift in space-use patterns in the season of 2019 compared to other years. This shift was particularly prominent when we compared the visit frequency in February, when ripe fruits of *S. spinosa* are usually most abundant. Guinea baboons visited the target area only once in 2019, whereas the visit frequency was 17 times on average in the years before and after the bushfires. The lower visit frequencies in 2020 may be the result of fire damage to *S. spinosa* trees in 2019 and, therefore, lower fruit productivity in the following year. The visit frequency to the target area in March 2019 was comparable to other years, which can be explained by the presence of other feeding trees in the area that bear ripe fruit in March (e.g., *Bombax costatum, Borassus akeassii, Piliostigma thonningii, Ficus ingens*). These trees are much taller than *S. spinosa* and therefore less likely to be impacted by bushfires as heavily as *S. spinosa*.

The pronounced change in space-use patterns indicates that Guinea baboons keep track of the spatio-temporal availability of a high-value food resource within their home range. It further shows the capacity to flexibly adjust space-use patterns and foraging decisions to changes in resource availability. Such flexibility may be particularly important for animals that live in environments with substantial seasonal variation associated with changes in food availability [56,57]. The behaviour is not simply habitual [21]. Instead, we suggest that certain environmental cues, including perhaps sunrise and sunset patterns, temperature, and the availability of specific food items [58], provide cues to the potential overall food availability in the target area. Once the animals have encountered the resources in the area and received the corresponding food rewards, they continue to visit the area until the resources are depleted. In 2019, the surprising absence of the fruits and the resulting prediction error prompted them to update their assessment of the area instantaneously [25]. In doing so, the animals likely reassessed and overwrote the informational value of potential environmental cues normally used to predict food availability. It is unlikely that Guinea baboons relied on olfactory or visual cues by the fruits when navigating towards the target area, as the sleeping sites of our study parties are predominantly located in the gallery forests alongside the Gambia River [24, 59], and the average approach distance from the sleeping trees to the target area was over 2000 m.

In addition, the Guinea baboons in the Niokolo-Koba National Park in Senegal travelled more linearly, faster, and started earlier towards the target area when *S. spinosa* was in season compared to when it was not. The linearity of travel was comparable to findings in other primate species (spider monkeys *Ateles geoffroyi yucatanensis* [37] chacma baboons [19]; western gorillas [6]; chimpanzees [36,60]). Travel speed towards the target area increased by about 14%, indicating a high motivation and anticipation of study parties to arrive at this high-value food resource. This notion is further supported by earlier times of departure from sleeping sites on days when moving towards the target area. The relative time of departure from the sleeping sites was also influenced by the minimum temperature, with later departures occurring at lower minimum temperatures, likely due to thermoregulatory activities (sunbathing) on cold mornings.

Similar patterns have been observed in other primate species. Chacma baboons approached valuable resources within their home ranges in a fast and linear fashion [19]. They left sleeping sites earlier when visiting fig trees, a highly preferred food resource, and only returned to less preferred seeds that are harder to ingest in the afternoons. Noser & Byrne [19] suggested that this behaviour may be a strategy to optimise the cost and benefits of foraging on a distant depletable, high-value food resource. Similarly, chimpanzee females left their nest sites earlier when breakfasting on preferred ephemeral fruit located far from their nesting sites [8], and skywalker gibbons (*Hoolock tianxing*) left their sleeping trees earlier when breakfasting on fruit instead of leaves [38], most likely to compensate for longer travel times to distant feeding sites as well as to mitigate interspecific competition.

From the available data, we cannot formally infer any specific navigational strategies. Euclidean maps assume highly detailed knowledge of locations and the ability to navigate between them by computing (novel) distances and angles, similar to a coordinate system or a coordinate-based map [20,61,62]. Topological or route-based maps, in contrast, allow for efficient navigation via the use of a set of interconnected, learned travel paths among relevant locations, often facilitated by using visual landmarks in the environment [5,20]. From long-term observations of our study parties, we know they consistently use “bottlenecks” in the landscape (e.g., paths to cross water streams, animal paths, and roads). Space use patterns, particularly at these highly conserved features of the landscape, strongly resemble route-based navigation, as in other baboon species [7,63,64].

In summary, our study provides strong evidence that Guinea baboons keep track of the spatio-temporal availability of a high-value food resource within their home range and flexibly adjust their space-use in response to unexpected changes. Rather than following fixed routines, they appear to rapidly update prior experience with current environmental information, suggesting a capacity for adaptive, goal-directed decision-making in a dynamic seasonal environment.

## Author contributions

Funding acquisition: D.Z., J.F.; conceptualisation: L.O., D.Z., J.F.; investigation: L.O.; data curation: L.O., R.M.; formal analysis: L.O., R.M.; visualisation: L.O., R.M.; writing – original draft: L.O., R.M., D.Z., J.F.; writing - review and editing: L.O., R.M., D.Z., J.F., supervision: D.Z., J.F.

## Data Availability

All data and scripts for analyses are available from the corresponding author upon request.

## Acknowledgements

We are grateful to the Direction des Parcs Nationaux (DPN) and the Ministère de l’Environnement et de la Protéction de la Nature (MEPN) de la République du Sénégal for the approval to conduct this study in the Niokolo-Koba National Park. We thank all past and present conservateurs of the park for their support throughout the years. We are grateful to all past and present CRP Simenti staff and field assistants for their support in the field and contribution to the collection of long-term GPS data, with special thanks to Djibril Coly, Amadou Bamba Diedhiou, and Chérif Younousse Kéba Camara for their outstanding efforts. We owe special thanks to Irene Gutiérrez Díez for her help in tirelessly mapping the target area.

## Ethics

Approval and research permission were granted by the Direction des Parcs Nationaux and the Ministere de l’Environnement et de la Protection de la Nature de la Republique du Senegal (protocol d’accord from 22 April 2019). Research was conducted within the regulations set by Senegalese agencies as well as by the Animal Care Committee at the German Primate Center (Göttingen).

